# Codon language embeddings provide strong signals for protein engineering

**DOI:** 10.1101/2022.12.15.519894

**Authors:** Carlos Outeiral, Charlotte M. Deane

**Affiliations:** Department of Statistics, University of Oxford, 24-29 St Giles’, Oxford, OX1 3LB, Oxfordshire, United Kingdom; Division of Biologics, Exscientia, Ltd., Oxford Science Park, The Schrödinger Building, Oxford, OX4 4GE, Oxfordshire, United Kingdom

**Keywords:** language models, protein representations, deep learning, protein engineering

## Abstract

Protein representations from deep language models have yielded state-of-the-art performance across many tasks in computational protein engineering. In recent years, progress has primarily focused on parameter count, with recent models’ capacities surpassing the size of the very datasets they were trained on. Here, we propose an alternative direction. We show that large language models trained on codons, instead of amino acid sequences, provide high-quality representations that outperform comparable state-of-the-art models across a variety of tasks. In some tasks, like species recognition, prediction of protein and transcript abundance, or melting point estimation, we show that a language model trained on codons outperforms every other published protein language model, including some that contain over 50 times more parameters. These results suggest that, in addition to commonly studied scale and model complexity, the information content of biological data provides an orthogonal direction to improve the power of machine learning in biology.

## 1 Introduction

Pretrained language models have become indispensable tools across many areas of computational protein engineering [1]. Most labeled protein datasets have limited size, therefore vast deep neural networks are first pretrained on a large, unlabelled corpus of sequence information, such as UniRef [2], with a self-supervised reconstruction objective. Self-supervised training endows the latent variables of the model with highly informative features, known as *learned representations*, which can then be leveraged in downstream tasks where limited training data is available. Learned protein representations are currently central to the state-of-the-art tools for predicting variant fitness [3–6], protein function [7, 8], subcellular localisation [9], solubility [10], binding sites [11], signal peptides [12], post-translational modifications [13], intrinsic disorder [14], and others [15, 16], and they have shown promise in the path towards accurate alignment-free protein structure prediction [17–21]. Improving learned representations is therefore a potential path to deliver consistent, substantial improvements across computational protein engineering.

Pathways towards more informative representations have hitherto followed two main directions. Methods have pursued the paradigm of *augmented scale*, where increasing model capacity monotonically increases performance [22]. While initial language models reached tens of millions [23] or hundreds of millions [24] of parameters, later developments have seen models with over 5 billion weights [19, 25, 26] with parameter counts exceeding the size of the training set. Improvements to *model architecture* have also consistently delivered performance gains. For example, the use of the T5 architecture in ProtTrans displayed consistent improvements in performance over the basic BERT model [8, 26]. The state-of-the-art fitness prediction method, Tranception, modifies the attention mechanism to explicitly attend to contiguous sequences of amino acids [6], increasing robustness and performance on deep mutational scanning benchmarks. Both directions are costly in human and computer time, require significant optimization, and appear to provide diminishing (logarithmic) returns.

An alternative pathway to improve learned representations may be to use biological data containing richer signals. While protein language models have so far focused on amino acid sequences, there is additional information contained in the DNA sequence encoding the protein. The language of protein-coding DNA relies on 64 nucleotide triads, known as *codons*, each of which encodes a specific amino acid or the end of a sequence. Although this 64-codon alphabet is highly degenerate, with most amino acids being encoded by up to six different codons, current research suggests that codons encoding the same amino acid (*synonymous*) are not used interchangeably. Synonymous codon usage has been correlated with protein structural features [27, 28], and nearly 60 synonymous mutations have been linked to human disease [29]. A recent experiment suggested that most synonymous mutations in yeast are strongly deleterious [30], although these results have since been contested [31, 32]. Codon usage has also been linked to protein folding, with ample evidence that changes in the codon sequence affect folding dynamics [33–36], the folding pathway [37] and even the amount of correctly folded protein [38]. This evidence suggest that synonymous codon usage contains valuable biological information, which could be exploited by machine learning models to enhance the signal-to-noise ratio in predictive tasks.

In this work, we demonstrate that pretraining a protein language model on codon sequences, rather than amino acid sequences, leads to informative protein representations that capture crucial biochemical characteristics. We examine the predictive power of these representations in a number of sequence-level prediction tasks, observing that these representations are comparable to, or superior to amino acid representations from similarly sized models. In several tasks, we observe that codon-based representations outperform all published state-of-the-art amino acid representations, including those from models with over 50 times more parameters. We conclude that finding more biologically informative representations of the data is a meaningful direction towards progress in deep protein engineering that does not suffer from the computational onerousness of larger scale, and is significantly simpler than – but also complementary to – improved model architectures.

## 2 Results

We developed a protein language model trained on protein-coding DNA (cDNA) and examined its ability to produce high-quality representations of protein sequences. We relied on the fact that the codon space is surjective, but not injective, to the amino acid space, therefore the former contains an amount of information higher or, at worst, equal to the latter (Figure 1b). To test this hypothesis, we trained a large language model with 86M parameters on a dataset of 9M non-redundant and diverse cDNA sequences identified from whole-genome sequencing (see Figure 1c). We refer to this model as CaLM (Codon adaptation Language Model). The training set was constructed from the European Nucleotide Archive [39], with significant preprocessing to limit redundancy and save computational cost. We also established a heldout data set consisting of representative sequences from seven model organisms across the tree of life. Details of model architecture, training protocol and dataset preprocessing are given in the Methods section.

**Fig. 1.**
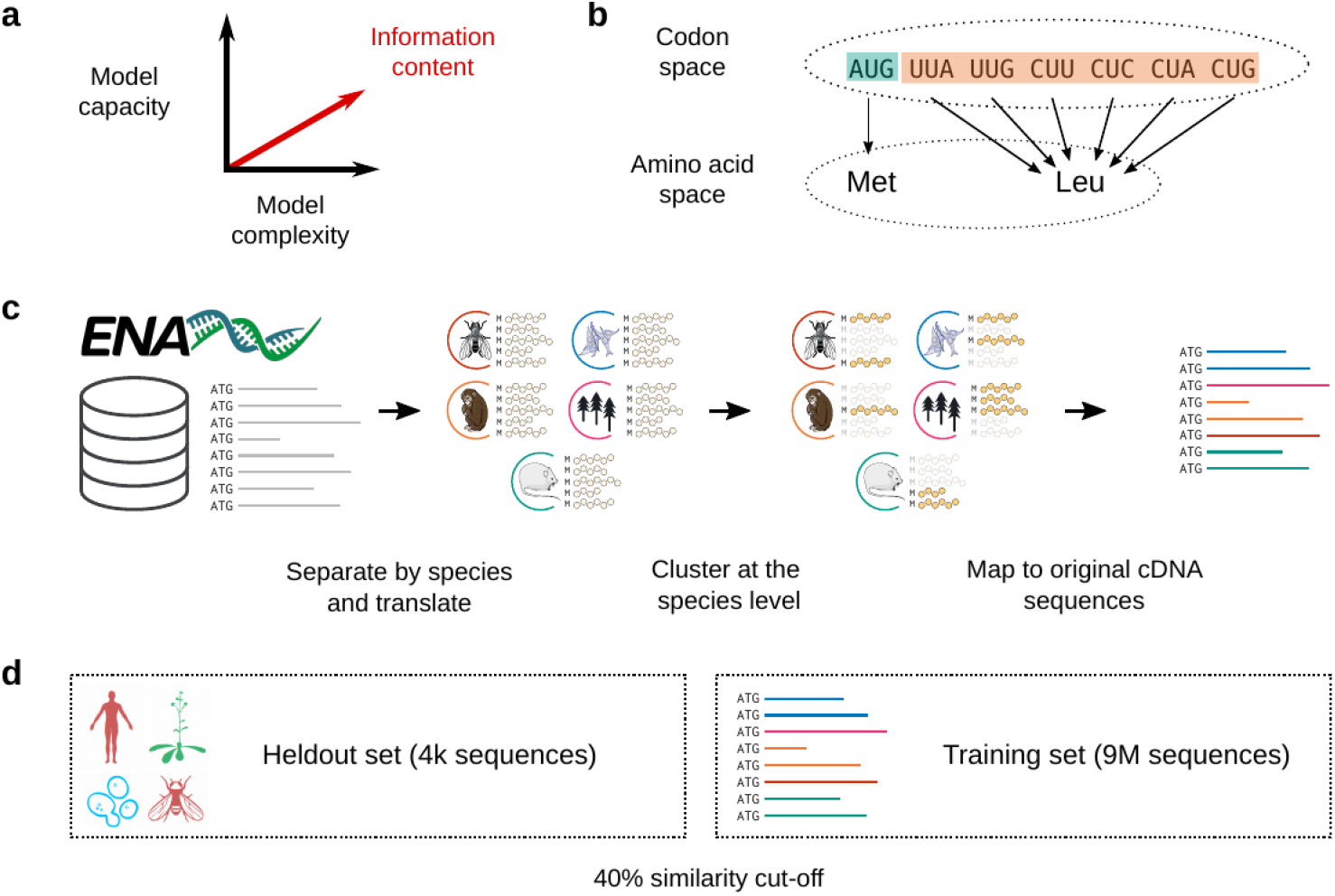
(a) Current research suggests that model performance may be improved by either increasing the number of parameters, or improving the architecture of the model. In this work, we propose a third, orthogonal dimension: the use of data with higher information content, in this case the codon, rather than the amino acid sequence. (b) The map between the codon alphabet and the amino acid alphabet is surjective, but not injective, hence there is more information in the codon space. (c) Processing of the training data. The original database of 114M cDNA sequences was divided into species and clustered at the protein level. (d) Scheme of the training and heldout datasets. As heldout, we selected 4,358 sequences from seven organisms spanning all kingdoms of life, and removed any sequence with 40% sequence identity or more from the training set.

### 2.1 Codon language models capture the biology of the genetic code

We first considered whether the learned representations from the codon language model captures the biochemistry underlying the genetic code. A model that has extracted meaningful representations should recognise the similarity of codons in the same wobble pair, a non-trivial task as the model represents individual codons as integers, with no features indicating nucleotide composition. The embedding should also capture the similarity of codons encoding amino acids with similar chemical behaviour, as do amino acids language models [24]. We tested these hypotheses by examining the embedding layer in CaLM (see Figure 2b). Dimensionality reduction shows that amino acids with similar behaviour cluster in similar regions of space. Clustering captures biochemical features that are not directly obvious from class labels: for example, the codons encoding alanine (“hydrophobic”) appear close to glycine (“special”), which reflects the small side chain both amino acids display. We also observed that pairs of codons that encode the same amino acid, or that are in the same wobble pair, are closer in space that others (*p <* 0.05, permutation test *N* = 10^7^).

**Fig. 2.**
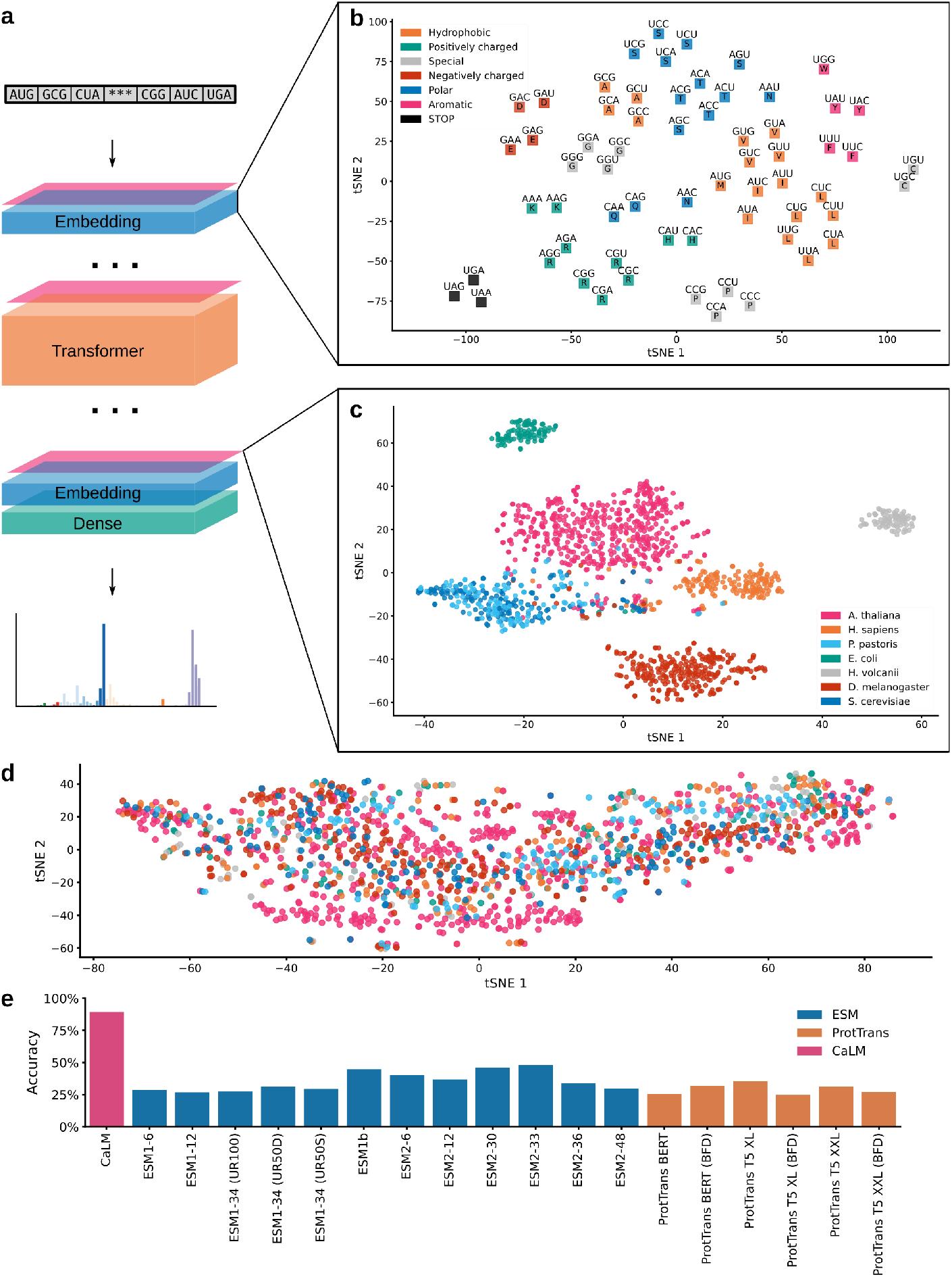
(a) Architecture of the Codon adaptation Language Model (CaLM). The sequence of codons is mapped to a continuous space via a trainable embedding, and passed through 12 layers of transformer encoders and a dense layer. The embedding is reversed at the end of the architecture. (b) Structure of the learned embedding space. Codons with similar biochemical properties (as shown by the colours) tend to occupy adjacent regions of space. Codons encoding for the same residue (amino acid single letter codes shown over the points) tend to be closer (*p* = 0.0169, permutation test *N* = 10^7^), as do codons in the same wobble group (*p* = 0.0203, permutation test *N* = 10^7^). (c) Structure of the latent space shown on one third of the sequences in our heldout dataset. The latent representations are distributed by species. (d) The embedding of the sequences in b using ESM2 [19], showing a lack of structure; see also Figure A3. (e) Accuracy of a nearest-cluster-center classifier at predicting the species of a sequence of the remaining two thirds of the heldout. The codon language model is significantly better than any other model (*p <* 10^*−*5^, Welch’s *t* test).

We then considered sequence representations of different organisms. In Figure 2c we display the embeddings of a third of the heldout dataset, which contains sequences with at most 40% sequence identity to any sequence in the training set. The sequences of prokaryotes *E*.*coli* and *H. volcanii* are significantly separated from their eukaryotic counterparts (*p* smaller than numerical precision, Welch’s *t* test). The sequences of *S. cerevisiae* and *P. pastoris*, which belong to the same order, appear intermixed. The region at the center of the plot where sequences from multiple organisms converge is enriched in highly conserved sequences such as ribosomal proteins or enzymes involved in the cell cycle. We also controlled for artifacts of dimensionality reduction by varying parameters and testing multiple algorithms (see Figures A1 and A2). We compared this clustered structure with representations from amino acid language models (see Figures 2d and A3), observing a less clear clustering. These findings suggest that codon representations capture richer sequence-level information that is not accessible to amino acid sequences alone.

We then tested the ability of the representations to assign cDNA sequences to species, using a simplified *k*-nearest centers classifier. Class centers were defined using one third of the heldout set, and tested on the remaining two thirds; the results are shown in Figure 2e. We observe that CaLM’s classification accuracy is almost twice as high as the best amino acid classifiers, and significantly superior (*p <* 10^*−*5^, Welch’s *t* test). Since our model is at the cDNA level, we controlled for the differential GC content across different species [40], observing that a logistic regression classifier would only achieve 48% accuracy, comparable to the predictions of amino acid representations. These observations suggest that the codon representations capture features of differential codon usage across distinct organisms that are not evident in the amino acid sequence.

Taken together, our results suggest that the codon language model can access biological features that are inaccessible to amino acid language models.

### 2.2 Codon language models match state-of-the-art performance on automated annotation tasks

We next examined whether the additional information contained in codon sequences can be used to improve protein engineering. Several benchmarks of language model representations have been proposed, such as TAPE [23], FLIP [41] and PROBE [8]. However, these datasets contain only amino acid sequences, and due to the loss of information, mapping amino acid sequences back to codon sequences is far from trivial. We therefore consider the performance of protein language models in four sequence annotation tasks where it was possible to recover the original codon sequence (see Methods): melting point prediction, solubility prediction, subcellular localization prediction and function prediction. Performance is assessed by 5-fold cross-validation (see Figure 3) after clustering the datasets with tight sequence identity cut-offs (dependent on the dataset, but ranging between 20% and 50%) to ensure removal of homologous sequences.

**Fig. 3.**
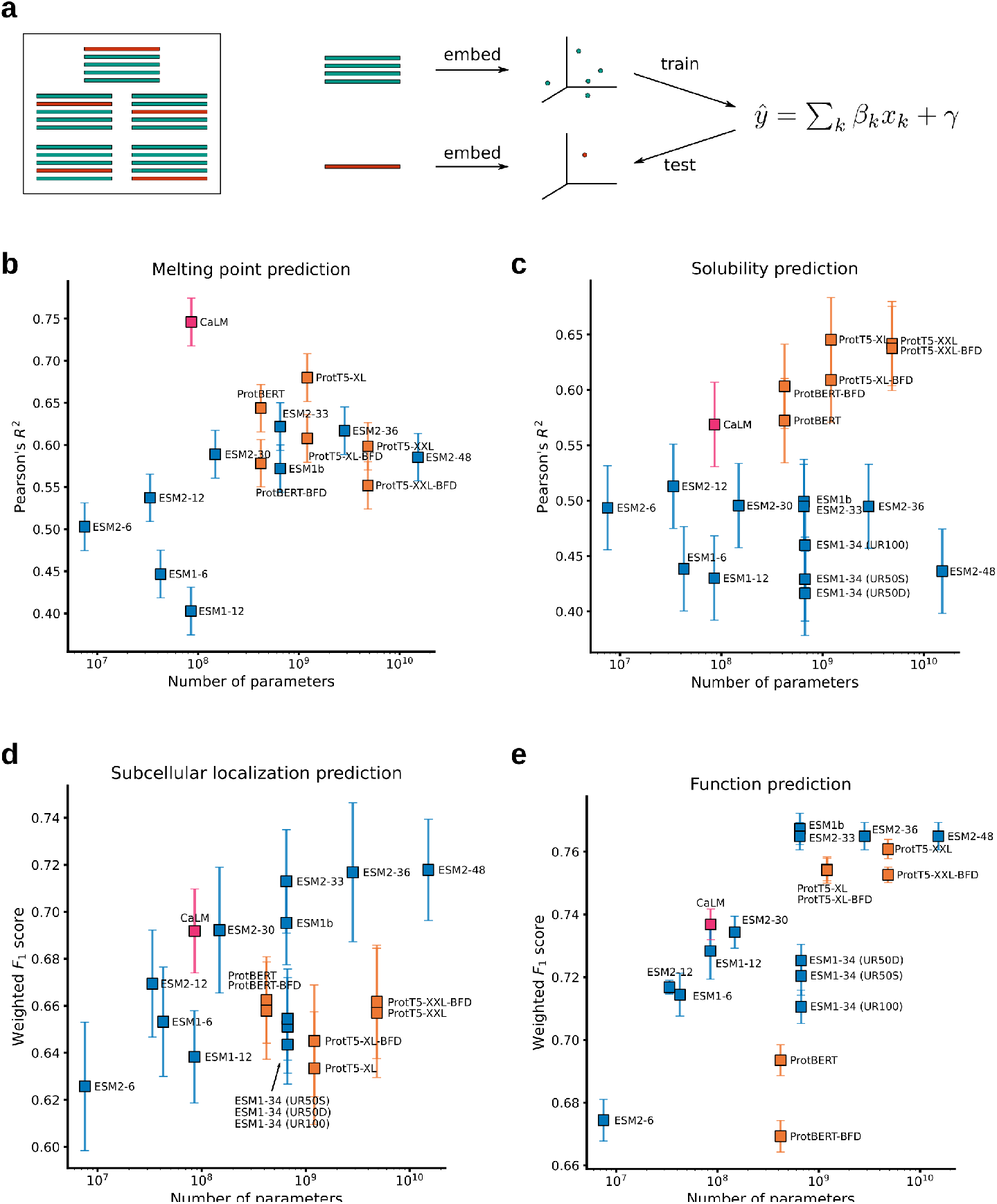
Assessment of codon language models in automatic annotation tasks. (a) Scheme of the 5-fold cross-validation protocol. The data is first divided in five groups which do not have any sequences with more than 40% amino acid identity. In every iteration, a model (a linear or logistic regression) is trained on four groups (green) and tested on the fifth group)red. The performance of the model with respect to the task and the number of parameters is subsequently shown for (b) melting point prediction, (c) solubility prediction, (d) subcellular localization classification and (e) function (Gene Ontology term) classification.

We observe that CaLM outperforms every amino acid language model of similar size across all tasks, and in some cases, also amino acid language models with over 50 times more parameters. In melting point prediction, CaLM achieves a Pearson’s *R*^2^ of 0.75, which is significantly better than any other method in the dataset. In solubility prediction, CaLM outperforms every model of the ESM family with which it shares architecture, and is comparable to the smaller models of the ProtTrans family, which are one order of magnitude larger and trained on two orders of magnitude more data. In subcellular localization and function prediction, the model outperforms all similarly sized architectures and is competitive many models of greater size and complexity.

We considered the hypothesis that the model may be relying on other information rather than the codon sequence. For example, the model may be learning stability information from species-level signals in the data. Many archaeal proteins are thought to be more stable due to the abundance of ion pairs in their structures [42], and since CaLM embeddings can accurately identify the source species of a protein, it might indirectly be using this information in prediction. We controlled for this hypothesis by comparing against a classifier including species information. We observed that CaLM’s predictive power increased from *R*^2^ = 0.74 to *R*^2^ = 0.78, and that while the absolute difference with the second best method narrowed from six to four percentage points, it was still significantly better (*p* = 10^*−*3^, Welch’s *t* test). The codon model thus demonstrates superior performance across various unrelated tasks, and against a variety of benchmarks.

We then considered the question of whether the improvement in prediction is due to synonymous codon usage. If patterns of codon usage contain valuable information, then performance should decrease if codon usage is somehow corrupted. We designed an experiment where the results in Figure 3b were repeated under the same conditions, but randomly mutating a fraction of the codons of both training and test datasets to other codons encoding the same amino acid (synonymous mutations). The results are shown in Figure A5. We observe that Pearson’s *R*^2^ drops from 0.75 to nearly half its value, 0.39, as the sequence of codons is fully randomised, a value that corresponds to the worst performance in the benchmark. These results suggest that the model is extracting useful information from the pattern of synonymous codon usage that is not available from the amino acid sequence.

These findings lead us to conclude that the codon sequence contains valuable information about protein properties that a codon language model is able to extract usefully.

### 2.3 Codon language models successfully capture features of omics datasets

As the codon language model presents improved performance in some tasks, arguably due to enhanced biological information, we then considered whether this approach can be applied to other tasks where codon usage is more important. Protein abundance inside the cell is one such task, as codon usage is well-known to present characteristic signatures in housekeeping genes [43]. To test this hypothesis, we constructed datasets of transcript (for the seven organisms) and protein abundance (for five organisms, due to data availability) in our heldout dataset, and evaluated the ability of CaLM to recover this information using 5-fold cross-validation. We also compared all amino acid-level models, as shown in Figure 4.

**Fig. 4.**
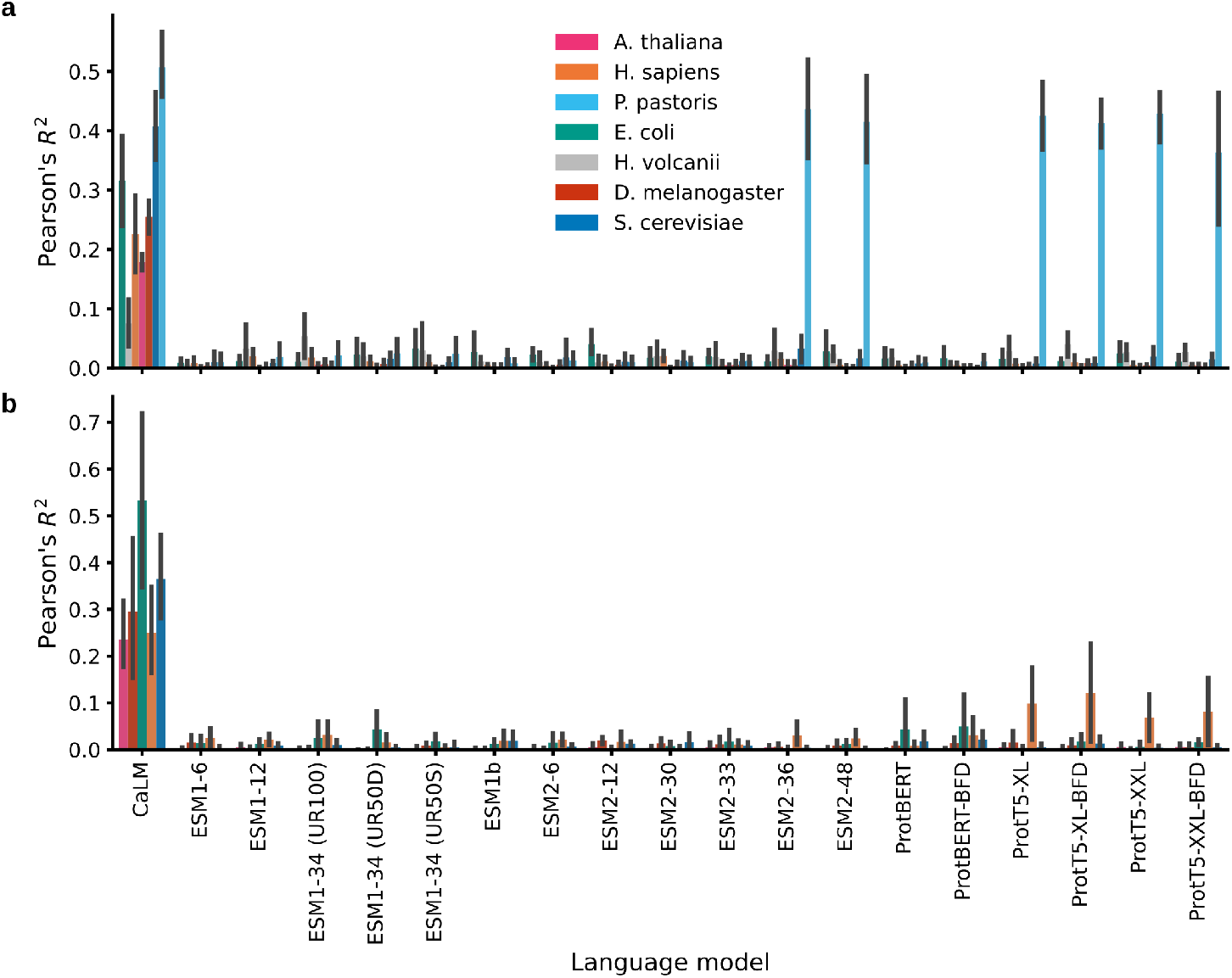
Assessment of codon language models at predicting the results of omics datasets using 5-fold cross-validation. (a) Transcript abundance prediction results for the seven organisms represented in our heldout dataset. The codon language model outperforms every other model for every organism, with the exception of *P. pastoris*, for which some of the 5 billion parameter models display similar performance. (b) Protein abundance prediction results on five of the seven organisms in our dataset, which were represented in the PAXdb repository [44]. The codon lanugage model outperforms every other model for every organism.

We observed that predictions from the codon model yield Pearson’s *R*^2^ in the 0.2-0.5 range, whereas most amino acid models fail to reach values of 0.1 for most species. One exception is the performance of transcript abundance in *P. pastoris*, where the largest amino acid language models (those with 3 billion parameters and higher) achieve moderate performance, although still worse than CaLM. One possible reason is the abundance of endogeneous signal peptides for secretion in *P. pastoris* [45], which may strengthen the performance of some amino acid models. In every case, the performance of CaLM is superior to the second best model (*p <* 0.005, Welch’s *t* test). This result further reinforces the hypothesis that codon language models are able to capture biological features of the sequences that are inaccessible to amino acid language models.

## 3 Discussion

In this work, we have shown that protein representations trained on codons, rather than amino acid sequences, exhibit significant advantage across a variety of downstream tasks. We find that our 86M parameter language model outperforms every other model of similar capacity, and in many cases, even models with over 50 times more parameters. We have provided evidence that this performance is due to the codon language model’s ability to capture patterns of codon usage across DNA sequences, and that this advantage disappears when codon usage information is corrupted.

Training models on cDNA comes at a negligible extra training cost, and appears to increase performance on all sequence-level tasks considered. Since high-throughput protein sequencing is done almost exclusively by translation of DNA sequences, the original coding sequences are publically available and can be used for training, although they have not been subject to the same standards of processing and annotation as protein sequence databases like UniRef [2]. We suggest that using cDNA, instead of simply amino acid sequences, to train protein language models, poses a clear pathway towards improving computational protein engineering.

Codon language models may also provide valuable evolutionary signals for alignment-free protein structure prediction, particularly in methods like ESMfold [19] and OmegaFold [18] that rely on language models to predict relationships between parts of the protein. Models based on cDNA may recover wider evolutionary relationships, such as synonymous mutations, which are evident at the nucleotide level but not at the amino acid level. Synonymous codon usage is known to relate to structural features [27, 28], and the connection between codon usage and protein folding [33, 36] may provide valuable signals to methods which are known to not capture the physics of folding [46]. We suggest that incorporating codon language models in the pipelines of alignment-free protein structure prediction may well provide a route with negligible cost towards accelerating high-accuracy protein structure prediction. We propose two main directions towards further improvements in protein representation quality. One is increased scale. The results in this paper have employed a simple model with only 86 million parameters, a size that pales in comparison to the standard model size in the latest publications. The dataset employed is also relatively small: merely 9 million sequences, in comparison to the 125 million used in the ESM family of models [19, 24] or the nearly half a billion in some ProtTrans models [26]. There exists a clear pathway towards improving representation quality by training billion-parameter models on datasets comprising hundreds of millions of DNA sequences.

The other potential direction for improvement is the development of multimodal models combining amino acid and coding sequences. Our ablation experiment showed that, in the absence of codon usage information, model performance decays significantly, to the point that it is inferior to every amino acid model in our dataset. However, since the model indirectly has access to the amino acid sequence, it should in principle have access to the same information as amino acid-only models. This divergence may be due to the lack of amino acid-level signals during training, so training models that combine amino acid and codon sequences could improve overall model performance.

Our results suggest that, concomitantly with advances in computational power and model architecture, leveraging richer biological data provides a clear direction towards improving the power of machine learning in biology.

## 4 Methods

### 4.1 Datasets

#### 4.1.1 Training and test data

We downloaded the coding sequences (CDS) of all organisms available in the European Nucleotide Archive with a timestamp of April 2022 (114,214,475 sequences). We considered only high-quality sequences pertaining to assembled genomes (data code ‘CON’). We filtered this dataset to remove all sequences with unknown nucleotides (symbols ‘N’, ‘Y’, ‘R’, and others), with a start codon different to ATG, containing interstitial stop codons, or where the number of nucleotides was not a multiple of three. To reduce redundancy while maintaining a representative dataset of codon variation across the tree of life, we grouped the entries by organism, translated the cDNA to protein sequences, and clustered the sequences of every organism at 40% amino acid identity using CD-HIT [47]. After backmapping clustered sequences to cDNA, the full dataset consisted of 9,858,385 cDNA sequences.

To enable rigorous testing of the model capabilities, we built independent heldout set containing sequences of seven model organisms spanning the tree of life: three eukaryotic multicellular organisms (*Arabidopsis thaliana, Drosophila melanogaster* and *Homo sapiens*), two eukaryotic unicellular organisms (*Saccharomyces cerevisiae* and *Pichia pastoris*), a bacteria (*Escherichia coli*) and an archaea (*Haloferax volcanii*). We queried GenBank for all cDNA sequences of every model organism according to the highest-quality assembly available, clustered them at 40% amino acid identity, and sampled 7.5% of the clustered sequences using random sampling stratified by protein abundance. Since no proteomic data was available for all organisms, we used transcript abundance measured by RNA-seq as a proxy for protein abundance (see Table A1 for data sources). To minimise the overlap between training and heldout set, we used BLAST to identify and remove homologous training set sequences with 40% cDNA sequence identity or higher to any sequence in the heldout set. After removing homologous sequences, the training set consisted of 8,771,938 sequences, and the heldout of 4,358 sequences.

#### 4.1.2 Evaluation datasets

To test the quality of the representations, we constructed several datasets to test the predictive performance of the learned representations. These datasets overlap with many published benchmarks of learned protein representations. With the exception of the transcriptomics dataset, where the sequence of codons can be inferred from the transcript, all available datasets reported only amino acid sequences. To obtain codon information, we mapped UniProt IDs to ENA entries using UniProtKB [48] and ignored all entries without a match. We also removed all sequences with unknown nucleotides, containing interstitial stop codons, or where the number of nucleotides was not a multiple of three.

##### 4.1.2.1 Melting temperature

We built a melting temperature dataset using proteome-wide denaturalization experiments reported in the Meltome Atlas [49]. We used the same splits and homology removal protocol as Dallago *et al*. [41], where data was clustered at 20% sequence identity.

##### 4.1.2.2 Subcellular localization

We used the SwissProt localization dataset as processed by the authors of DeepLoc 2.0 [9]; this dataset is also part of the FLIP benchmark set [41]. We used the same clustering as the original authors. Although cluster sizes were slightly different due to UniProt IDs that could not be mapped, we noted that fold size variance was small enough to conserve the original splits.

##### 4.1.2.3 Solubility

We used the human solubility proteome profiling experiments by Sridharan *et al*. [50]. As a proxy for solubility, we used the average protein abundance determined in the SDS-treated fraction of the experiment. We clustered the sequences of at 40% amino acid identity using CD-HIT [47].

##### 4.1.2.4 Gene Ontology

We used the Gene Ontology dataset published by Unsal *et al*. [8], which used experimental annotations from UniProtKB/Swiss-Prot and UniProtGOA.

##### 4.1.2.5 Transcriptomics

We collected RNA-seq datasets for all seven model organisms from the Gene Expression Omnibus (GEO), the EMBL-EBI Expression Atlas, the primary literature and the Sequence Read Archive (SRA); data sources are reported in Table A1. The corresponding assemblies of all organisms are reported in Figure A2). We estimated transcript abundances of all proteins in the assembly in transcripts per million, and mapped these values to the sequences in the heldout dataset.

##### 4.1.2.6 Proteomics

The Protein Abundance Database (PAXdb) [44] was queried for protein abundance data on the seven model organisms used in this work. Samples for A. thaliana, D. melanogaster, E. coli, H. sapiens and S. cerevisiae. Dataset coverages were greater than 95% for all five organisms except for A. thaliana, with 76% coverage. This data was used to assign protein abundances to all proteins in the heldout dataset.

### 4.2 Model details

#### 4.2.1 Model architecture

CaLM is a language model inspired by the ESM family of architectures [19, 24] (see Figure 2a for an architectural diagram). The model consists of three blocks: a learnable embedding, a stack of transformer encoder layers, and a prediction head. The input sequence is a vector of T tokens, integers which each represent a codon or a special character. For example, the number 11 corresponds to the start codon “AUG”, whereas the number 68 represents a special character “⟨mask⟩” used for masking. The alphabet is composed of the 64 codons, plus five special characters: “⟨mask⟩” for masking, “⟨cls⟩” indicating the start of a sequence, “⟨eos⟩” to indicate the end of a sentence, “⟨pad⟩” for padding and “⟨unk⟩” for potentially unknown codons. No prior knowledge is given to the model: codons are represented in an abstract manner, and there is for example no way in which the model can learn that codons “AUG” (token number 11) and “AUA” (token number 8) differ only at a nucleotide level.

The vector of tokens, with dimensions [T] is mapped into a learnable latent space of dimension 768 by the embedding layer, leading to a vector of size [T, 768]. This vector is then passed through multiple layers of transformer encoders, following the architecture of Devlin *et al*. [51]. The transformer layers contain 12 attention heads, with dimension 768, and the feed-forward neural network part of the transformer has dimension 3,072. Following Rives *et al*., we use pre-normalisation to increase stability [24]. Since the multi-head attention layer is equivariant to permutations of the input tokens, we use Rotary Positional Embeddings (RoPE) [52] to enable learning of sequential features. The vector at the end of the transformer stack is referred to as a “representation”, and is also of size [T, 768]. The representation vector is the main focus of the paper, although the model is trained alongside a language head that predicts the probability of every token at a given position. The language head consists of a feed-forward neural network, followed by layer normalisation, and a product by the inverse learned embedding matrix. The output logits, when transformed via a softmax, provide an uncalibrated probability distribution over codons.

#### 4.2.2 Model training

We trained the model in a self-supervised manner using dynamic masking. In every training batch we masked 25% of the input tokens at random. Of the masked tokens, 80% were substituted by a special token “⟨mask⟩” indicating masking, 10% were substituted by another codon at random, and the remaining 10% were left untouched. We used the cross-entropy loss to train the model to predict the right token, considering only the 25% of the tokens that had been masked. Following Rives *et al*. [24], we used no dropout.

Sequences were trimmed to a maximum size of 1,024 tokens, a number that we found empirically to be sufficiently large to enable efficient learning while preserving computational efficiency. This is consistent with other published models, as 96% of all UniParc entries have fewer than 1,024 amino acids [24]. Sequences larger than 1,024 codons were subsampled at random at every batch. The size of all sequences in every batch was padded to the maximum sequence in the batch.

We trained the model using the AdamW optimizer with a learning rate of 1e-4 and default parameters otherwise. The learning rate was warmed up from 0 to 10^*−*4^ during the first 1,000 gradient steps, and subsequently decayed with a cosine function that reaches zero after 120,000 steps. Gradients were accumulated to an effective batch size of 1,000 examples, or approximately 256,000 tokens. To monitor training, 1% of the training set was reserved at random as validation. The model reported in this work was trained on 4 NVIDIA Quadro RTX4000 GPUs for 40 days (66,000 gradient steps, 14 full epochs). Training was manually stopped after observing no validation loss improvement after 8,000 steps.

### 4.3 Model evaluation

#### 4.3.1 Embedding visualization

We used the t-distributed Stochastic Neighbours Embedding (tSNE) method to reduce the dimensionality of token and sequence embeddings and enable visualization. We used the implementation of tSNE in sci-kit learn 0.23.2 [53] with default parameters. To ensure reproducibility, we performed sensitivity analysis on the perplexity hyperparameter, as well as comparisons to an alternative dimensionality reduction, Uniform Manifold Approximation and Projection (UMAP), which are reported in the Supplementary Information. Plots reported in the main text use the default values of the sci-kit learn implementation, as well as a maximum of 10,000 iterations to ensure convergence.

#### 4.3.2 Source prediction

Protein source prediction was benchmarked with a simple nearest-centroid algorithm. We divided the heldout dataset into two splits: parameter estimation (33%) and test (66%). Using the parameter estimation set, we computed the centroid of all sequences corresponding to a given species. At the test stage, we assigned a sequence to a species according to the centroid with the smallest L2 distance.

##### 4.3.2.1 Property prediction

We tested the models using 5-fold cross-validation. Splits were done using scikit learn 0.23.2 with default parameters and shuffling, except in the subcellular localization task where we used the splits published by DeepLoc.

## Acknowledgments

The authors would like to thank Dr. Oliver M. Crook and Tobias H. Olsen for enlightening discussions.

## Declarations

Carlos Outeiral thanks the United Kingdom’s Engineering and Physical Sciences Research Council for financial support through an EPSRC Doctoral Prize (EP/T517811/1) and a postdoctoral fellowship (EP/W522582/1).

## Appendix A Supplementary Information

**Table A1.**
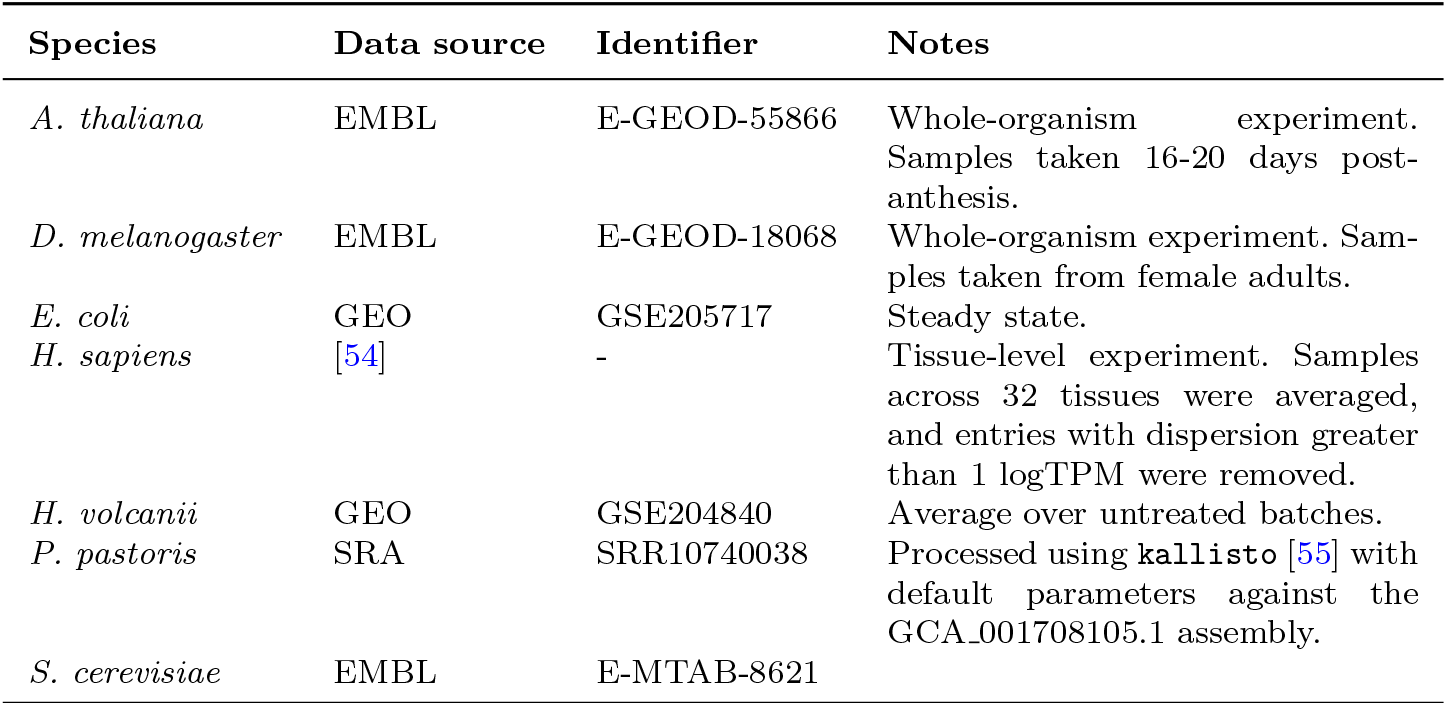
Transcriptomic dataset sources

**Table A2.** Assemblies

| Species | Data source | Identifier | Notes |
| --- | --- | --- | --- |
| *A. thaliana* | GenBank | GCA_000001735.1 |  |
| *D. melanogaster* | Ensembl | BDGP6.32 |  |
| *E. coli* | GenBank | GCA_000259695.1 |  |
| *H. sapiens* | Ensembl | GRCh38.107 |  |
| *H. volcanii* | GenBank | GCA_000025685.1 |  |
| *P. pastoris* | GenBank | GCA_001708105.1 |  |
| *S. cerevisiae* | GenBank | GCA_000146045.2 |  |

**Fig. A1.**
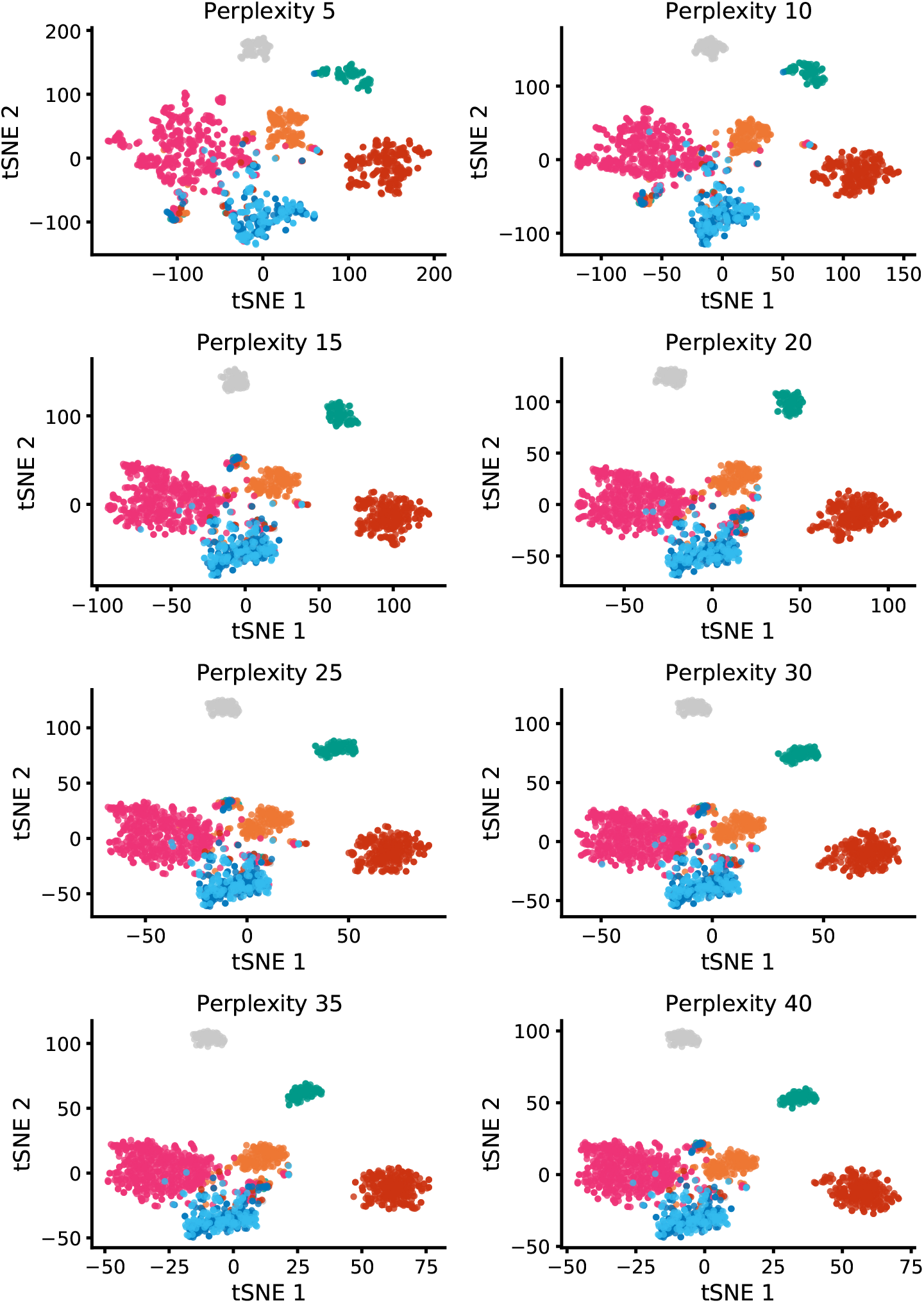
Comparison of the tSNE embedding presented in Figure 2c with different perplexity values.

**Fig. A2.**
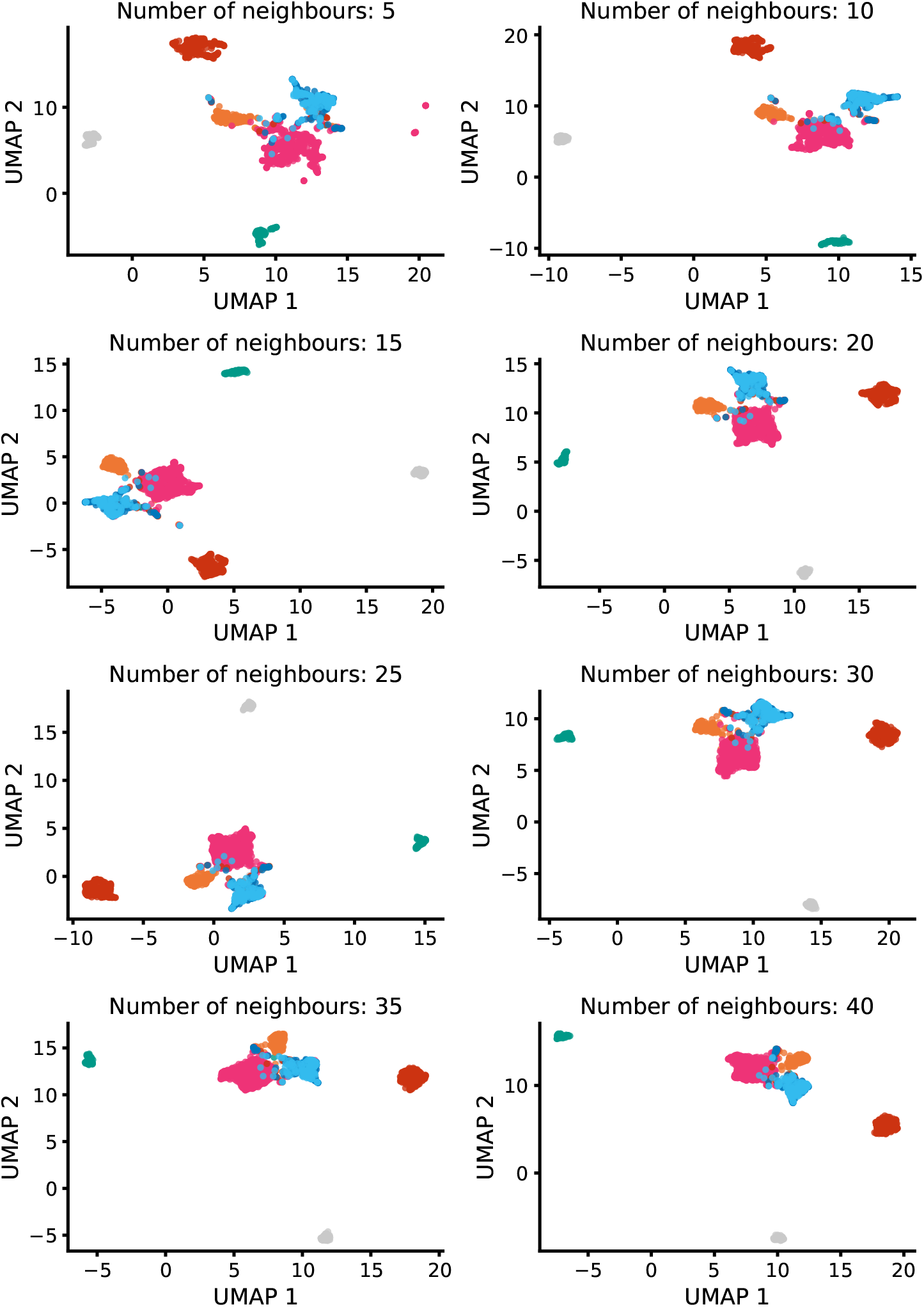
Comparison of the tSNE embedding presented in Figure 2c using UMAP and different numbers of neighbours.

**Fig. A3.**
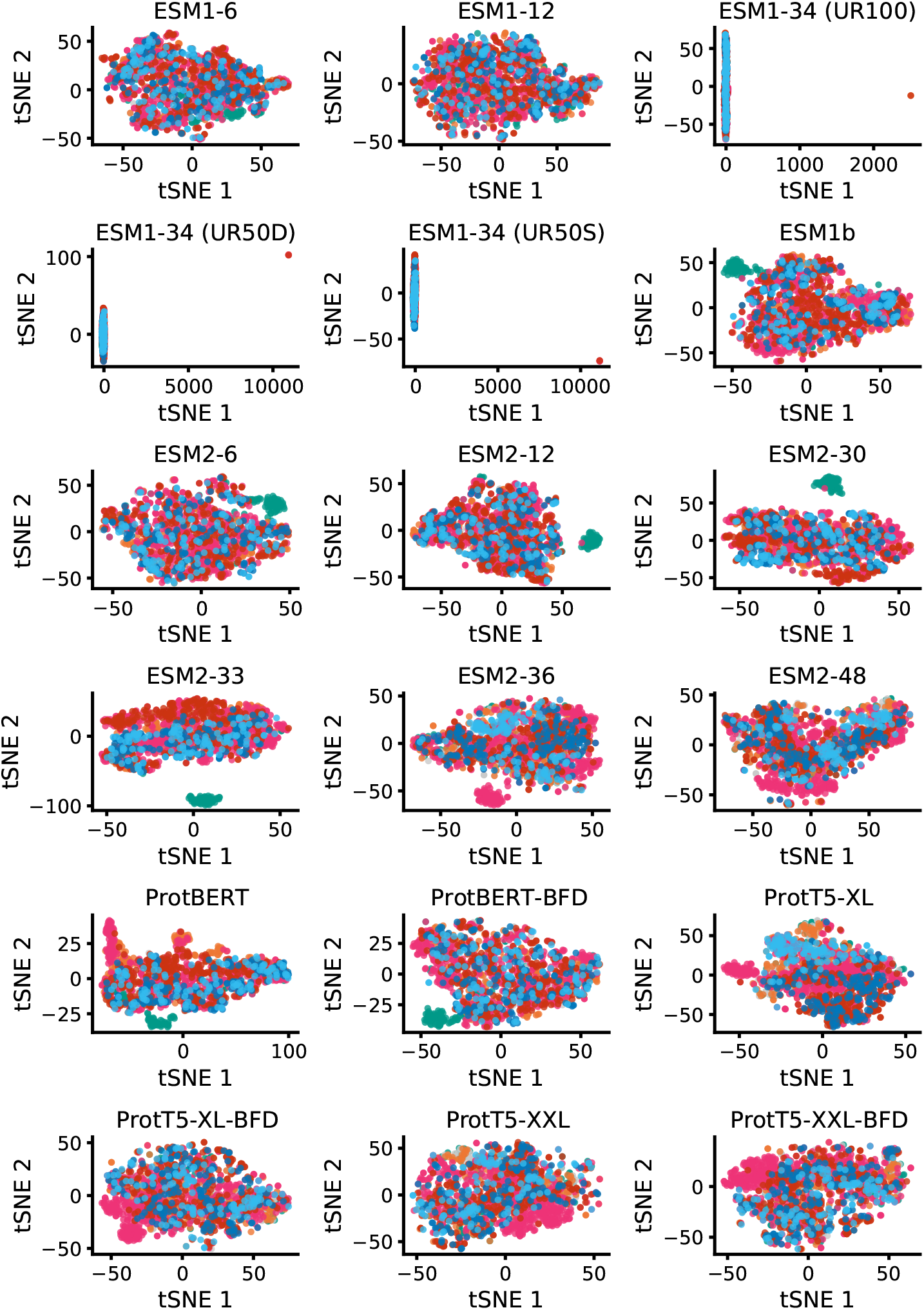
Replicates of the tSNE embedding presented in Figure 2c using different amino acid language models.

**Fig. A4.**
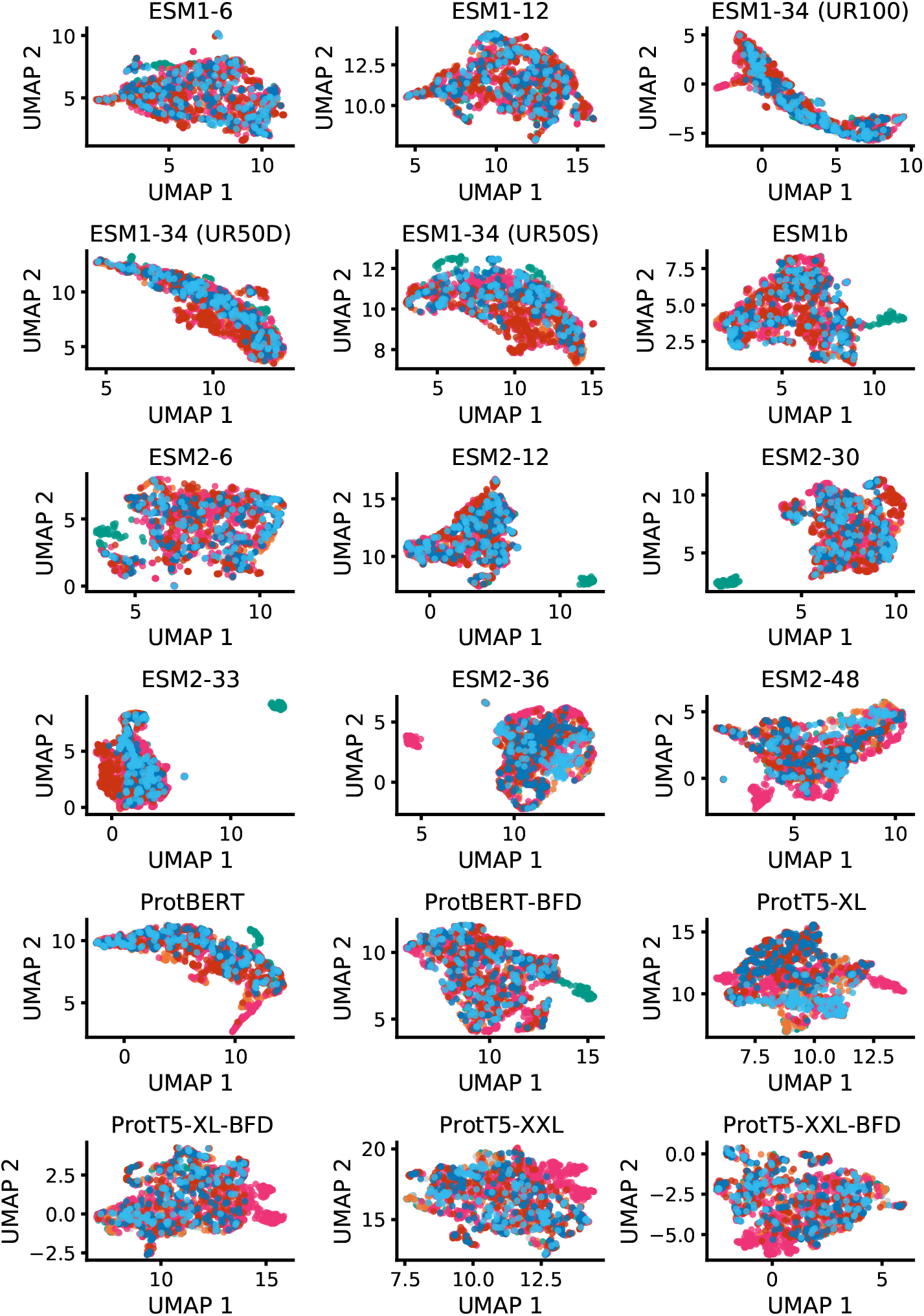
Replicates of the tSNE embedding presented in Figure 2c using UMAP and different amino acid language models.

**Fig. A5.**
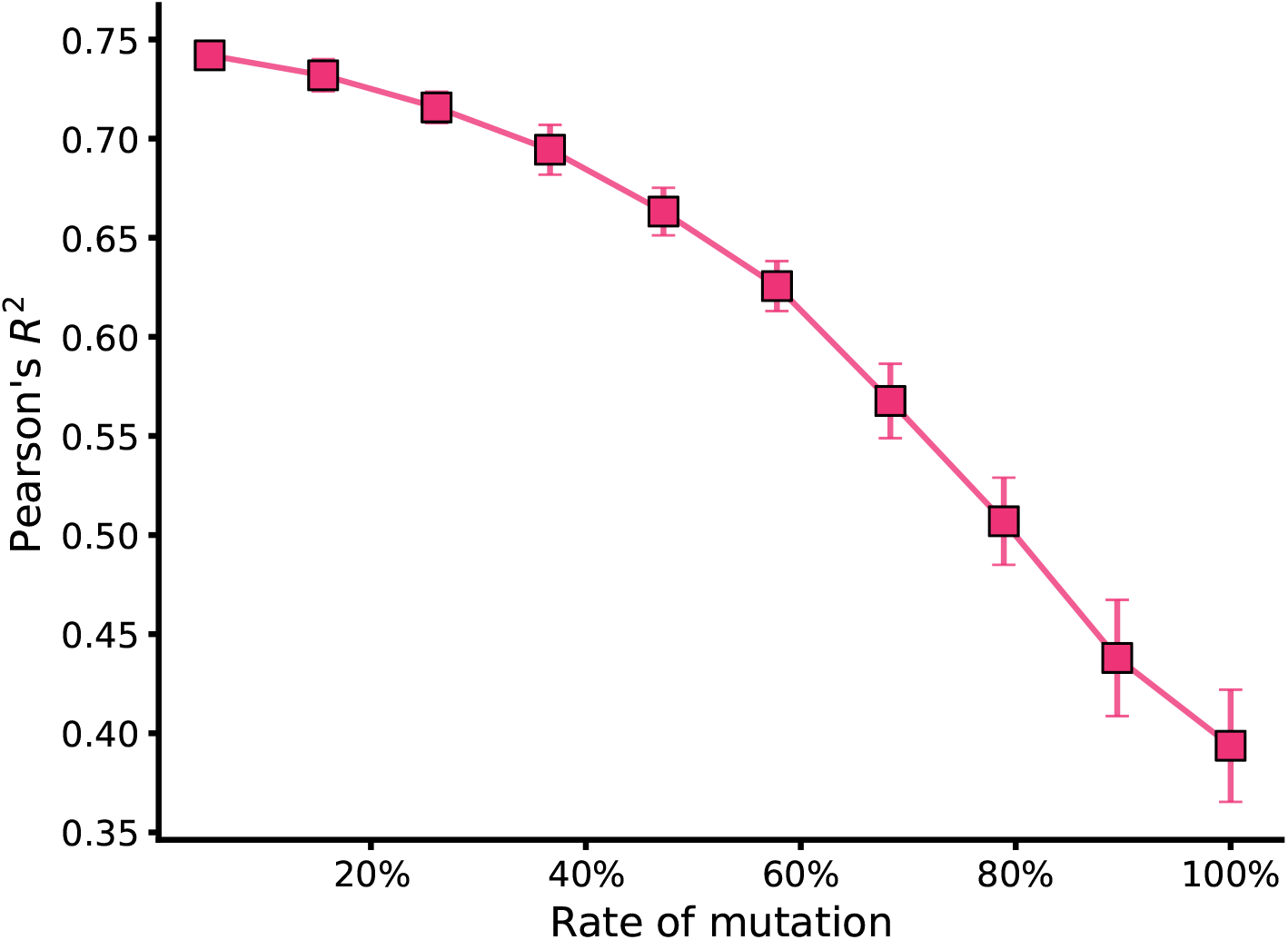
CaLM’s performance at predicting melting point with increasing rates of synonymous codon mutations. The correlation between predictions and ground truth values drops by nearly half as the rate of mutations approaches 100%, suggesting that codon usage information is fundamental for CaLM’s performance.

